# Preoptic galanin neuron activation is specific to courtship reproductive tactic in fish with two male morphs

**DOI:** 10.1101/515452

**Authors:** Joel A. Tripp, Isabella Salas-Allende, Andrea Makowski, Andrew H. Bass

## Abstract

Species exhibiting alternative reproductive tactics (ARTs) provide ideal models for investigating neural mechanisms underlying robust and consistent differences in social behavioral phenotypes between individuals within a single sex. Using phospho-S6 protein (pS6), a neural activity marker, we investigate the activation of galanin-expressing neurons in the preoptic area-anterior hypothalamus (POA-AH) during ARTs in midshipman fish (*Porichthys notatus*) that have two adult male morphs: type I’s that reproduce using an acoustic-dependent courtship tactic or a cuckolding tactic, and type II’s that only cuckold. The proportion of pS6-labelled galanin neurons increases during mating by courting type I males, but not cuckolders of either male morph or females, and is not explained by vocalization, eggs in the nest, or cuckolders present during mating. These differences within the same behavioral context indicate a male phenotype-specific role for galanin neurons in mating interactions, providing the most direct evidence to date for the role of a specific neuronal population in the differential activation of one mating behavior over the other in species exhibiting ARTs. More broadly, together with their known role in mammalian mating, the results suggest a deep-rooted, phylogenetically shared function for POA-AH galanin neurons in reproductive behavior. As such, these findings also provide new insights into the evolutionary relationship between POA-AH populations involved in social behavior regulation in teleosts, the most species-rich vertebrate group, with those in the more highly differentiated POA-AH of mammals.

**Significance Statement:** Galanin-expressing neurons in the preoptic area-anterior hypothalamus (POA-AH) are associated with mating and parental care in mammals. Here, we show that POA-AH galanin neurons are also active in a teleost fish during mating in a social context specific to one of two male morphs of a species with alternative reproductive tactics (ARTs). Together, the results suggest a key role for galanin-expressing neurons in the performance of reproductive-related social behaviors that is shared between distantly related vertebrate lineages and for galanin neuron activation in the evolution of ARTs. The results also help to clarify the relationship between molecularly-defined populations in the teleost POA-AH with the more highly differentiated mammalian POA-AH.

---

Studies of species exhibiting alternative reproductive tactics (ARTs) reveal how hormonal and neural mechansims can lead to the production of widely divergent social behaviors, ranging from territorial aggression and courtship to parental care, between individuals within the same sex (1, 2). ARTs have evolved numerous times among teleost fish (3). This includes the highly vocal plainfin midshipman (*Porichthys notatus*), a well-established neuroethological model with two male morphs (Fig. 1A). Type I males express either a courting reproductive phenotype or a cuckolding phenotype in which they steal fertilizations at the nests of courting males (4, 5), while type II males exclusively cuckold. Courting type I males alone excavate nests under rocks in the intertidal zone, produce a courtship vocalization known as a hum (Movie 1), provide parental care for developing eggs and larvae, and aggressively defend their nest and young from both type I and type II cuckolding males (6–8). The two male morphs further differ in developmental trajectory: type II males reach sexual maturity at a younger age, are smaller, and lack a hypertrophied vocal-motor system coupled to a suite of morphological, neurophysiological and hormonal traits (2, 6, 17, 9–16). Many of these type II traits are shared with females (Table S1). These sharp differences between male morphs, coupled with the behavioral flexibility of type I’s to either court or cuckold, provides an opportunity to explore how neural mechanisms of adult social behavior regulation are influenced by both developmental history and behavioral context.

**Figure 1.**
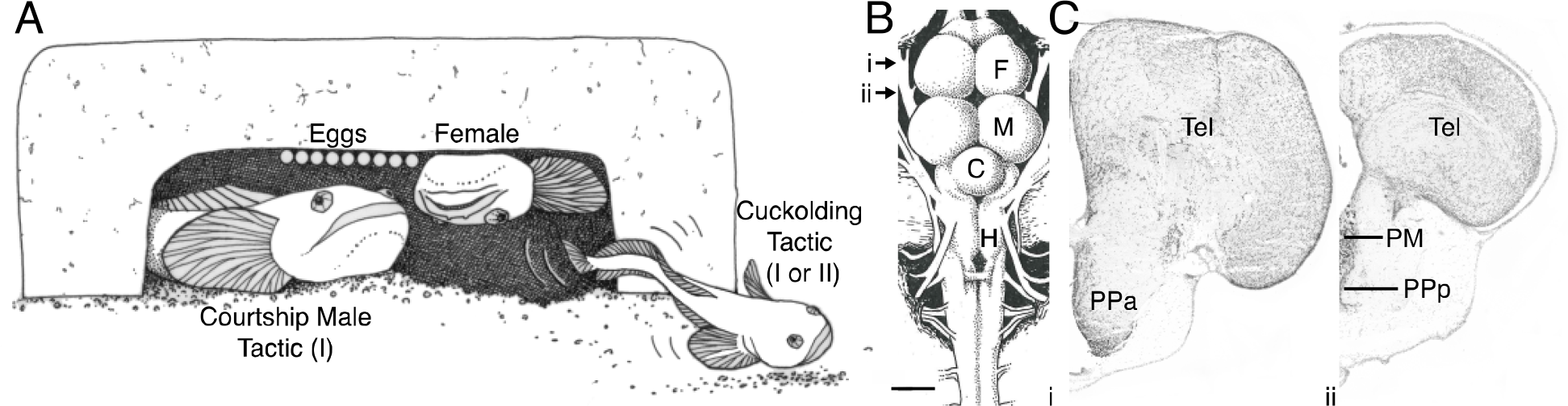
Midshipman reproductive behavior and preoptic area-anterior hypothalamus. A) Line drawing of midshipman nest during mating. Courting type I male in nest with female laying eggs, and satellite mating male outside fanning sperm into the nest. Both type I and type II males satellite mate. B) Overhead drawing of midshipman brain. Arrows indicate level of sections shown in Ci and Cii. Scale bar is 1 mm. C) Cresyl-stained coronal sections through midshipman brain at rostral (i) and caudal (ii) levels of the preoptic area. Images in Figs. 2-5 taken at similar level as (i). Abbreviations: C, cerebellum; F, forebrain; H, hindbrain; M, midbrain; PM, magnocellular preoptic area; PPa, anterior parvocellular preoptic area; PPp, posterior parvocellular preoptic area; Tel, telencephalon.

To date, no studies identify specific cell types that are differentially active during tactic-specific behaviors in teleosts or other vertebrate lineages that exhibit ARTs (3, 18–21). Putative candidates come from studies of neuropeptide-containing neurons in the preoptic area (POA)-anterior hypothalamus (AH), namely the oxytocin-vasopressin family of nonapeptides that has emerged as a key player in social behavior plasticity (22–26). This includes investigations showing inter- and intrasexual differences in the size and/or number of nonapeptide containing neurons in sex/tactic changing fish (23, 25), and nonapeptide actions on descending pathways from the POA-AH to the brainstem vocal network in midshipman (15).

Studies of neuropeptide regulation of social behavior in mammals have also traditionally focused on the nonapeptide family (27, 28). More recently, the neuropeptide galanin, which has established functions in pituitary hormone release and regulation of feeding (29–33), has received increased attention for its role as a regulator of social behavior (34, 35). Early studies of the POA-AH in mammals show a role for galanin in rat sexual behavior (36, 37) and identify populations of galanin neurons that is active during mating in male ferrets and mice (38, 39). The latest experiments demonstrate that galanin neurons promote parental care in mice of both sexes (39–41). High throughput RNA sequencing analyses also show higher galanin transcript expression in whole brain samples of parental (courting type I-like morph) than sneaker (cuckolding type II-like morph) male sunfish (*Lepomis macrochirus*) (42). More specifically, we showed for the POA-AH of midshipman that galanin transcript levels are elevated during mating in courting type I compared to cuckolding type II males, while cuckolding type I’s have intermediate expression levels (43). Though such associations are suggestive, it remains unknown whether galanin-expressing neurons play an active role in regulating ARTs during specific behaviors as in mammals.

The advent of transcriptomics along with molecular markers for neural activity offer new opportunities to investigate proximate mechanisms underlying behavioral variation in species lacking sequenced genomes (44). Guided by our transcriptomic study (see above), we use phosphorylated S6 protein (pS6), a marker of neural activity (45), to test the hypothesis that POA-AH galanin neurons (POA-AH^Gal^) are differentially activated during mating, but not other eproductive-related behaviors, in individuals exhibiting the courting type I male tactic and not in males of either morph exhibiting the cuckolding tactic or in females.

## Results

### POA-AH^Gal^ neurons

The POA-AH of midshipman, like other teleosts, includes three major divisions: anterior and posterior parvocellular nuclei flanking the anterior commissure and a magnocellular nucleus caudal to the commissure (Fig. 1B) (46, 47). Most POA-AH^Gal^ neurons were observed in the anterior parvocellular POA-AH, though some are in the posterior parvocellular and magnocellular POA-AH.

### POA-AH^Gal^ activity increases only in courting type I males during mating

Because we previously reported increased expression of galanin trancripts in the POA-AH of courting type I males compared to cuckolding type II males during mating (43), we predicted that POA-AH^Gal^ neurons are active only in courting type I males during mating. Additionally, based on prior studies of mammals, we predicted that there would be no increase in POA-AH^Gal^ neuron activation in females (38, 39). To test these hypotheses, we created a courting/cuckolding mating behavior paradigm where groups of type I and type II males were held in paired divided aquaria, with each chamber containing a single artificial nest to which gravid females could be introduced (Fig. 2A). A female was added to each chamber and when one female entered a nest, the other nest was blocked using a plastic mesh cage to prevent mating. Courting type I males (n=5), cuckolding type I males (n=5), cuckolding type II males (n=5), and females (n=3) were collected from nests where mating occurred along with paired control animals from neighboring nests that were prevented from mating over seven total spawning trials (Fig. 2A). Fewer females were collected, as some that remained gravid were kept to be reused in additional spawning experiments. Each mating animal and its paired control were collected two hours after they began mating (see Methods for rationale). Midshipman cuckoldry consists of either satellite mating in which a cuckolding male inserts its tail into the nest and fans sperm onto eggs (Movie 2), or sneak mating in which the cuckolder enters the nest, behaviorally mimics a female, and attempts to fertilize eggs as they are laid (5, 6). To reduce variation within groups, we collected only satellite type I and type II cuckolding males in this experiment, which were more frequently observed than sneaking males.

**Figure 2.**
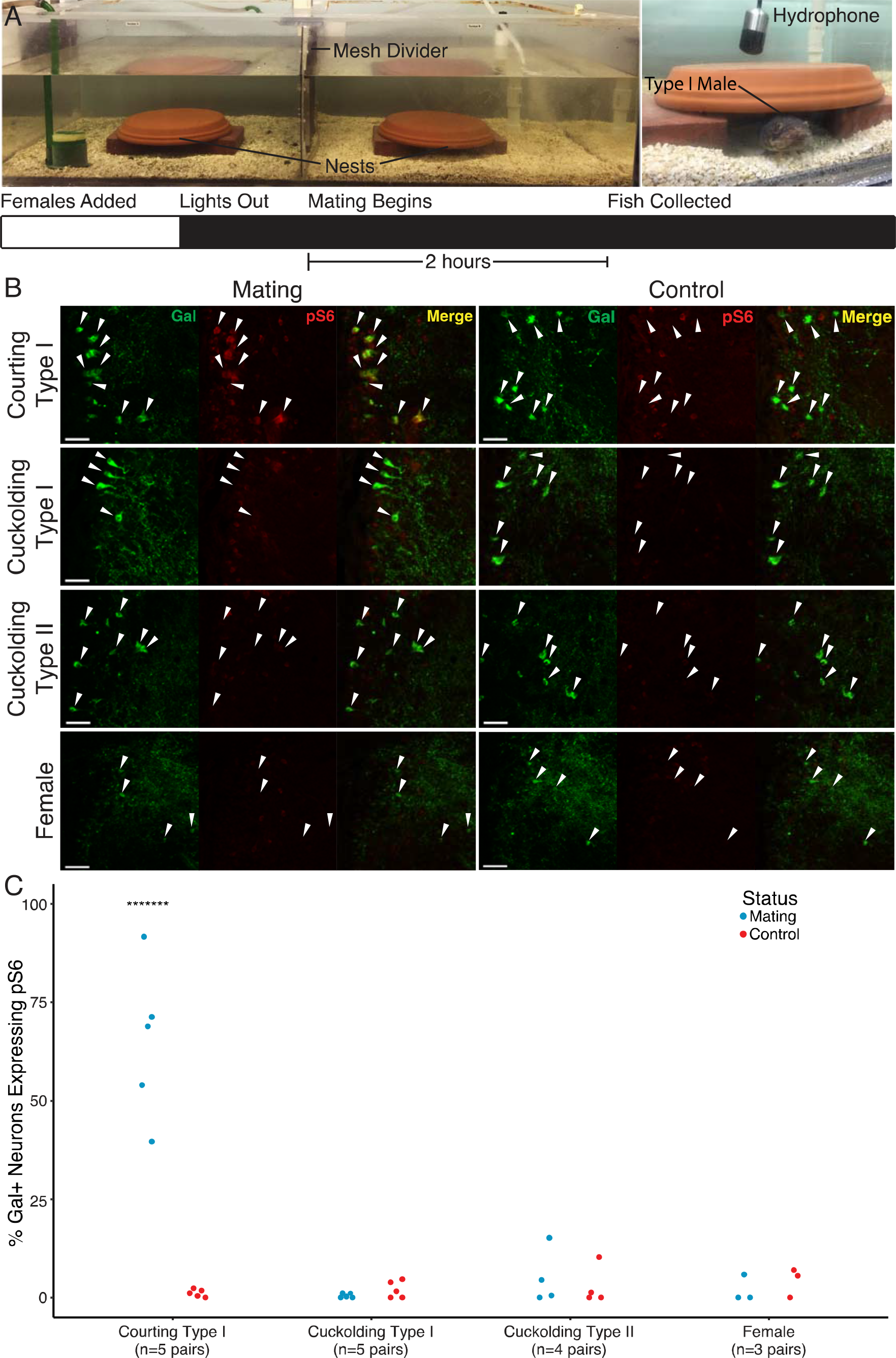
Mating behavioral experiment. A) Divided tanks used for mating experiment (left) and close up image of type I male in an artificial nest (right) with experiment timeline (below). Photographs taken during lights on to enable easy visualization of setup. B) pS6 expression in POA-AH^Gal^ neurons of mating (left) and non-mating controls (right) for courting type I, cuckolding type I, cuckolding type II males, and females (non-mating control courting type I alone inside nest covered by mesh cage; control cuckolding type I and II males and females blocked from accessing a nest). For each panel, left image shows galanin label in green, middle image shows pS6 label in red, with merged image on the right. White arrowheads indicate location of Gal cell bodies. Scale bars 50μm. C) Proportion of POA-AH^Gal^ neurons expressing pS6 in mating (blue) and control (red) courting type I, cuckolding type I, cuckolding type II males, and females. ANOVA F_(7,26)_=34.22, p=1.5*10^−11^. ^*******^p<1*10^−7^ Tukey’s test.

The brains of all animals in this experiment and those reported below were dissected immediately after collection, and prepared for immunohistochemistry (IHC) with tissue sections labelled by antibodies raised against galanin and pS6 (see Methods). One pair of type II male brains (mating and control) was damaged during IHC and was removed from further analyses. The proportion of POA-AH^Gal^ neurons expressing pS6 is increased in type I males engaged in the courting tactic, but not type I or type II males engaged in the cuckolding tactic, or mating females (Fig. 2B, C). ANOVA comparison shows a statistically significant difference among groups (F_(7,26)_=34.22, p=1.5*10^−11^). Tukey post-hoc comparison reveals that courting type I males engaged in mating have a significantly higher proportion POA-AH^Gal^ neurons expressing pS6 than all other mating and control groups (p<1.00*10^−7^), while there are no significant differences between all other mating and control groups (p>0.99 for all other comparisons).

The type I male courting tactic is a suite of several related behaviors that contribute to reproductive success (6). Because these behaviors can occur in quick succession (e.g., humming immediately precedes female entry to the nest), or may be interleaved with mating itself (e.g., bouts of aggression toward cuckolding males), it is possible that one of these component behaviors explains the POA-AH^Gal^ activation seen here in the courting male phenotype. To determine whether POA-AH^Gal^ activity is specific to mating, rather than other related component behaviors, we next tested whether these cells were similarly activated during type I male courtship humming, care for eggs, or nest defense against attempted cuckolders.

### POA-AH^Gal^ activity does not increase during courtship vocalization

Courtship humming immediately precedes mating and contributes to female nest localization and mating (6, 8, 48). Because there are increases in both galanin transcript expression in the POA-AH (43) and, as shown above, in POA-AH^Gal^ activity in courting type I males, galanin expression and neuron activity may have been related to humming specifically, but not mating itself. To test whether the observed POA-AH^Gal^ activation in type I males is associated with courtship humming, we recorded male vocalization using hydrophones, and collected eight males two hours after the onset of humming (Fig. 3A) along with eight non-humming control males housed in the same divided aquarium (see description above, Fig. 2A). There is no significant increase in the proportion of POA-AH^Gal^ neurons expressing pS6 in type I males humming in the absence of females compared to non-humming control males (t_(7.2434)_=−1.8073, p=0.1122, Welch two sample t-test, Fig. 3B, C). Additionally, for humming males, pS6 expression in POA-AH^Gal^ neurons is not significantly correlated with time spent humming during the two hours following hum onset (p=0.3021, r=−0.418533, Fig. 3D), or with the latency between hum offset and collection from the nest (p=0.2451, r=0.465508, Fig. S1).

**Figure 3.**
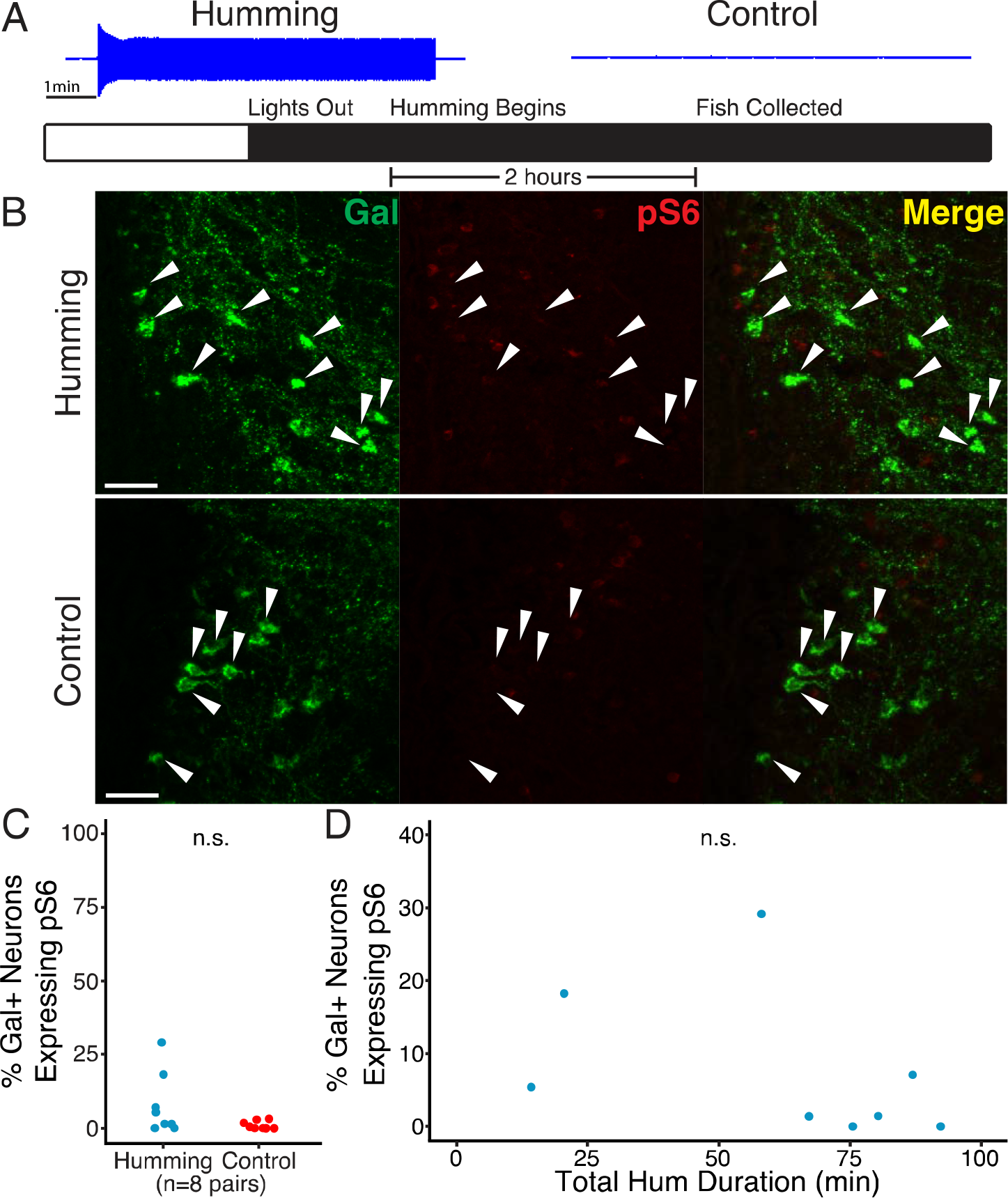
Courtship humming behavioral experiment. A) Example of recording from a humming male (left) and a non-humming male (right) and timeline of experiment. B) pS6 expression in POA-AH^Gal^ neurons of humming and non-humming control animals. For each panel, left image shows Gal label in green, middle image shows pS6 label in red, with merged imaged on the right. White arrowheads indicate location Gal cell bodies. Scale bars 50μm. C) Proportion of POA-AH^Gal^ neurons expressing pS6 in humming (blue) and non-humming control (red) type I males (t_(7.2434)_=−1.8073, p=0.1122). D) Relationship between pS6 expression in POA-AH^Gal^ neurons and total hum duration during experiment for humming males (p=0.3021, r=−0.418533). n.s, not significant.

### POA-AH^Gal^ activity does not increase during egg care

Immediately after mating, females depart from the nest and type I males begin egg care which includes fanning and brushing of eggs, and protecting them from predators (6, 7, 49, 50). To test whether the observed POA-AH^Gal^ activation is associated with egg care behavior that begins following mating, we collected singly housed type I males from nests containing eggs (n= 7), or control nests without eggs (n= 6) two hours after onset of the dark period, when midshipman are most active (51) (Fig. 4A). Observation of videos taken during the egg care experiment show that four of seven males given nests with eggs performed both fanning and brushing during the two hours before sacrifice. Additionally, two males were observed brushing but not fanning, and only one male did neither during the experiment. The proportion of POA-AH^Gal^ neurons expressing pS6 in type I males in nests with eggs is not significantly increased compared to males in nests without eggs (t_(11)_=0.66927, p=0.5171, two sample t-test, Fig. 4B-C), and there was no significant correlation between pS6 expression in POA-AH^Gal^ neurons and bouts of egg brushing, defined as putting mouth to eggs (p=0.6706, r=−0.1978782, Fig S2A), or with amount of time spent fanning fins (p=0.06185, r=0.7312215, Fig. S2B). This result was surprising, given the importance of medial POA-AH galanin neurons in mouse parental care (39, 40). However, we cannot rule out a role for POA-AH^Gal^ neurons in midshipman parental care, possibly during later stages of larval development (see Discussion).

**Figure 4.**
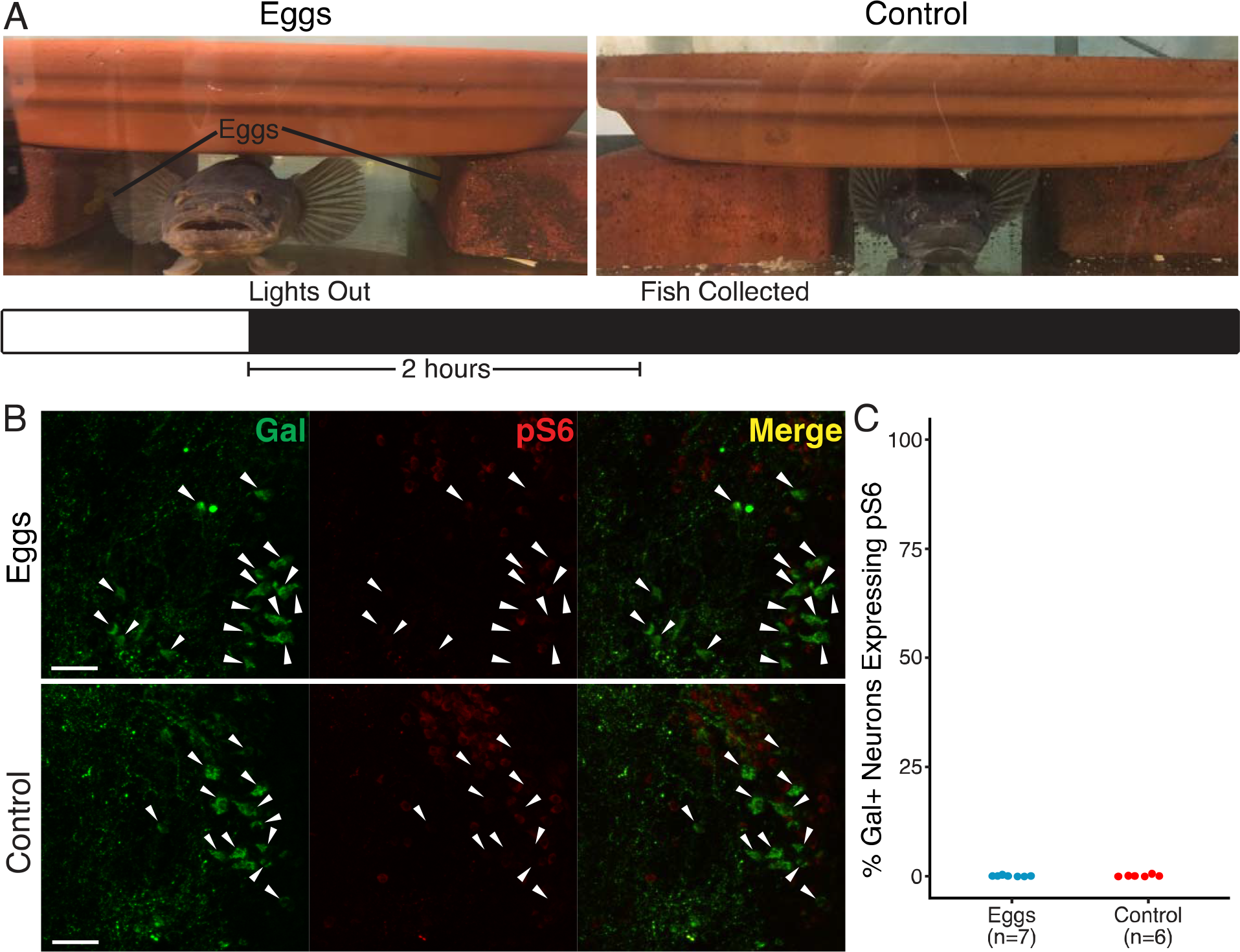
Egg care behavioral experiment. A) Experiment design. Example nests with eggs (left) and control nest without eggs (right) and timeline of experiment (below). Photographs taken during lights on to enable easy visualization of setup. B) pS6 expression in POA-AH^Gal^ neurons of males in nests with eggs and control animals without eggs. For each panel, left image shows Gal label in green, middle image shows pS6 label in red, with merged imaged on the right. White arrowheads indicate location of galanin cell bodies. Scale bars 50μm C) Proportion of POA-AH^Gal^ neurons expressing pS6 in males in nests with eggs (blue) and control nests without eggs (red). Independent t-test t_(11)_=0.66927, p=0.5171.

### POA-AH^Gal^ activation does not require territory defense

Courting males in our courting/cuckolding mating paradigm engage in territorial defense against cuckolders in addition to courtship towards and mating with a female (5, 6). To determine if the increased POA-AH^Gal^ activity observed during type I male mating was due to nest defense against cuckolders, we conducted another mating experiment in which courting type I males mate either with, or without potential cuckolders present in the aquarium. We collected courting type I males mating with and without cuckolders present two hours after a female entered and remained in the nest (Fig. 5A). Because laboratory animals were leaving reproductive condition, and we were no longer able to hand collect reproductive males due to the lack of low tides at the end of the breeding season (6, 52), we were only able to obtain n=2 mating males for each group in this experiment. Due to this small sample size, which would result in low power in any statistical test, we use only descriptive statistics for pS6 expression in these males. For both males collected during mating with cuckolders, video analysis confirms cuckolding attempts by other males and aggressive behaviors by courting males during the two-hour experiment. However, because we recorded using red light, we were not able to record behavior within the nest without potentially disrupting behavior, so it is possible that some aggression towards cuckolders was missed and we do not attempt to correlate aggressive behaviors with pS6 expression.

**Figure 5.**
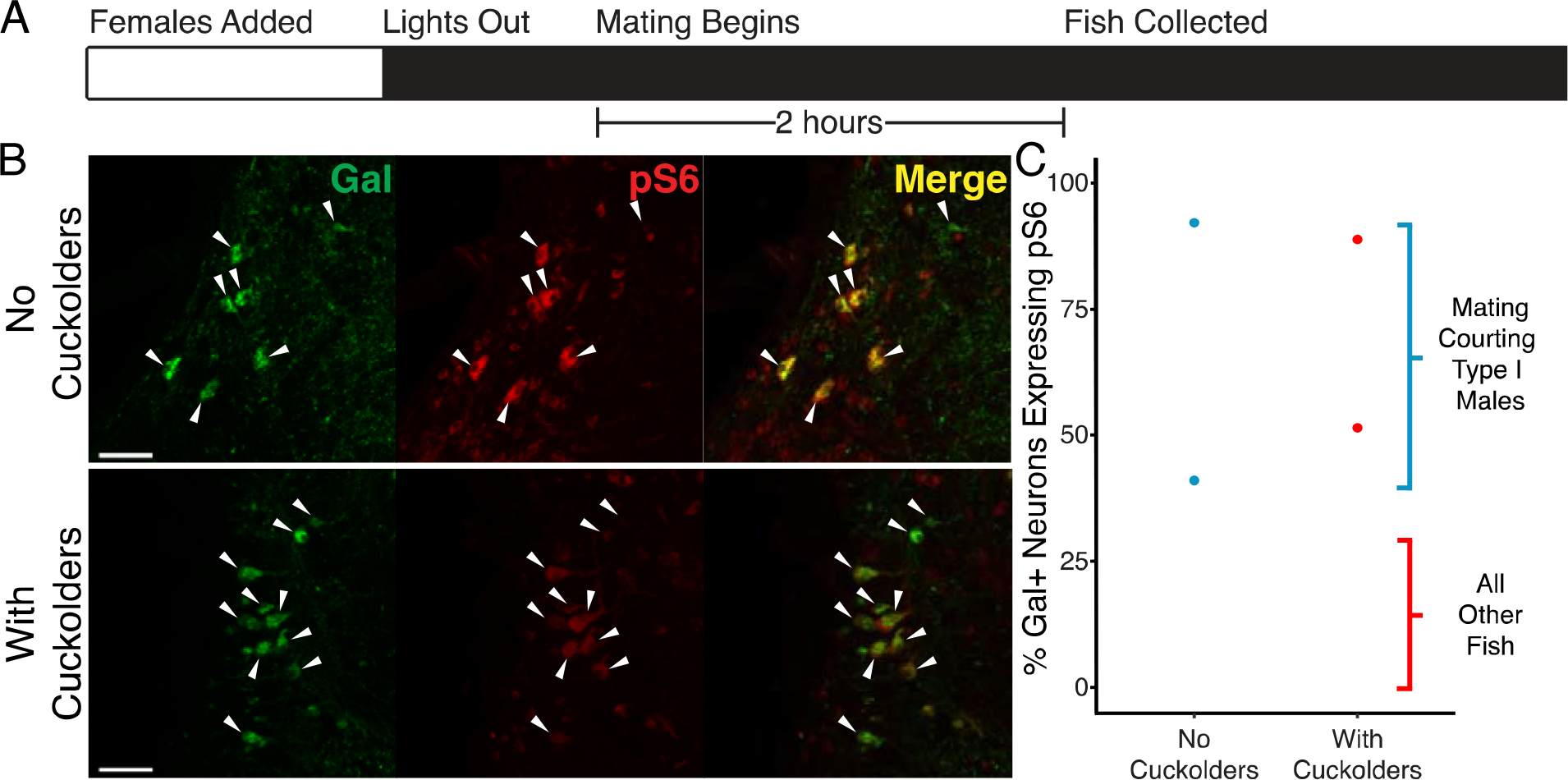
Nest defense behavioral experiment. A) Timeline of experiment. B) pS6 expression in POA-AH^Gal^ neurons of males mating with no cuckolders and with cuckolders present. For each panel, left image shows Gal label in green, middle image shows pS6 label in red, with merged imaged on the right. White arrowheads indicate location of Gal cell bodies. Scale bars 50μm. C) Proportion of POA-AH^Gal^ neurons expressing pS6 in males mating without cuckolders (blue points) and with cuckolders present (red points). Blue shaded region indicates the range of POA-H^Gal^ neurons expressing pS6 in courting type I males that mated in the earlier experiment (Fig. 2C). The red shaded region indicates the range of POA-AH^Gal^ neurons expressing pS6 in all other animals in all other experiments (Fig. 2C, Fig. 3C, Fig. 4C).

Both groups had similar levels of pS6 expression in POA-AH^Gal^ neurons (Fig. 5B, C). Courting type I males mating without cuckolders present have pS6 expression in 66.78±36.93% (mean±standard deviation) of POA-AH^Gal^ neurons, while males mating with cuckolders present show pS6 expression in 70.44±26.90% of POA-AH^Gal^ neurons. Strikingly, for each of the mating males in this experiment, the proportion of POA-AH^Gal^ neurons expressing pS6 (Fig. 5C) is within essentially the same range as that reported in the first experiment (Fig. 2C) for courting type I males, each of which had at least one cuckolder at their nest during mating. One male that mated without cuckolders present has a highly similar, but 1% greater proportion of POA-AH^Gal^ neurons expressing pS6 than the maximum expression seen in the first experiment. While the possibility remains that there are subtle differences in POA-AH^Gal^ activity between males defending against cuckolders and males without cuckolders present, our results indicate that the presence of cuckolders during mating is not necessary for increased POA-AH^Gal^ neuron activity.

## Discussion

Understanding the role of neuropeptide-containing neurons in the POA-AH remains critically important for establishing the mechanisms underlying individual variation in social behavior both within and between species of vertebrates. Our experiments take advantage of the expression of male ARTs in a well-studied neuroethological model to investigate the behavioral function of a specific POA-AH cell type, galanin-expressing neurons. As explained in more detail below, the results are significant for several reasons. First, although prior studies of gene expression in the brain of teleosts (including our own of the midshipman POA-AH) suggest a role for galanin in male mating behavior, the current investigation is the first to test whether galanin-expressing neurons are active during teleost mating. In this regard, the current results provide essential behavioral tests of hypotheses generated by transcriptomic research (42, 43, 53). More broadly, the results allow us to identify intrasexual differences in the activity of these neurons during reproductive behaviors that are central to the evolution of ARTs. This has made it possible to identify which behavioral modules comprising alternative tactics— in this case, acoustic courtship, mating, nest defense, egg care, cuckolding—are or are not coupled to galanin neuron activity. Second, by identifying a mating behavior role for POA-AH^Gal^ neurons in fish, we demonstrate a shared function for these neurons with distantly related mammals in the regulation of a specific reproductive behavior. Third, this functional information about POA-AH^Gal^ neurons elucidates the evolutionary relationship between specific cell types in teleost and mammalian POA-AH.

### Evolution of Alternative Reproductive Tactics in Teleosts

Galanin is a widely studied peptide (54), but relatively few experiments, so far limited to mammals, investigate the potential involvement of galanin-expressing neurons in regulating social behavior (38–40). Taken together, our current results show a role for a specific cell type, POA-AH^Gal^ neurons, in the courtship mating tactic of type I male midshipman fish. The results complement prior work showing elevated galanin transcript expression in the POA-AH of courting type I males during mating (43), in whole brain samples of reproductively active male cichlids (*Astatotilapia burtoni*) that lack ARTs (53), and in parental compared to sneaker male bluegill sunfish (*Lepomis macrochirus*) that utilize similar ARTs as midshipman, with parental and sneaker males being behavioral analogues of type I courting and type II males, respectively (42).

Despite the behavioral flexibility exhibited by type I male midshipman that can switch between courting and cuckolding dependent on social context (4), prior studies of neuroendocrine mechanisms focus on differences between the male morphs showing, for example, how vocalization could depend on the actions of different nonapeptides in the two morphs (15). The current investigation shows that the activation of a single neuropeptide-expressing population can vary both within and between the two male morphs. So, while our earlier studies emphasize the importance of developmental history in sculpting male morph-specific neuroendocrine and behavioral phenotypes, the present results demonstrate the importance of reproductive behavioral context, in this case the activation of one neuropeptide cell type during a single reproductive tactic (courtship) that is absent during the alternative tactic (cuckoldry), regardless of developmental morph. The differential activation of these neurons during alternative mating tactics points to a potential role of circuits including POA-AH^Gal^ neurons in the evolution of ARTs.

### Comparisons with Mammals

Our results are consistent with reports of increased expression of the immediate early gene c-fos in POA/POA-AH^Gal^ neurons following mating in male, but not female, mice and ferrets (38, 39), linking the neuroendocrine basis for differential regulation of individual, context-dependent reproductive-related behaviors in species with ARTs to those without ARTs such as rodents (see Introduction). The increase in POA-AH^Gal^ activity seen here in courting males may be related to interactions with the mating female at the entrance of and within the nest. These include blocking the entrance to prevent females from leaving once they enter, lateral pressing against the female prior to egg-laying, biting and maneuvering the female within the nest during egg-laying and sometimes forcing females into the nest (6). The precise role in relation to these more nuanced mating interaction behaviors remains unclear. For example, POA-AH^Gal^ neurons may be regulating the performance of these behaviors, or alternatively may be promoting mating interactions in general over other components of the courtship tactic (e.g. humming or nest defense). Because we used red light to observe behavior in this study, we were unable to illuminate and record behavior within the nest without potentially disrupting nocturnal courting male behavior.

Addtionally, our results inform recent comparisons between cell groups in the POA-AH of teleosts and mammals. Recent anatomical and developmental studies identify the POA of tetrapods as part of the subpallial telencephalon, separate from the hypothalamus (55, 56). A study of zebrafish (*Danio rerio*) recognizes the POA-AH as a distinct “morphogenetic entity” lying between the telencephalon and hypothalamus (57), perhaps reflective of it being an amalgam of neuronal groups comprising the POA and AH of mammals. The POA-AH of teleosts includes neurons synthesizing the nonapeptides isotocin (oxytocin homologue) and arginine-vasotocin (arginine vasopressin homologue) that are part of the AH of tetrapods; hence, our referring to this region in midshipman as the POA-AH (58, 59). Nonapeptide-synthesizing neurons within magnocellular and part of the parvocellular POA-AH in teleosts have led, in part, to their being proposed as homologues of the tetrapod paraventricular (magnocellular) and supraoptic (parvocellular) nuclei in the AH (59, 60). The predominant location of galanin-containing neurons active during male mating interactions in the anterior parvocellular POA-AH of midshipman supports its comparison to the medial preoptic nucleus of mammals that includes galanin-expressing neurons active during parental care (39).

The results of our egg care experiment were quite surprising, given the role of POA-AH^Gal^ neurons in mouse parental care (39, 40). We expected to see POA-AH^Gal^ activity related to egg care because courting type I males are the sole providers of parental care (6, 7), including care for eggs they did not fertilize after taking over a nest (61). More recent fiber photometry experiments in mice show that medial POA-AH^Gal^ neurons are only active during specific “pup-directed” parental care behaviors (40). However, we observe both fanning and brushing in males given nests with eggs and neither behavior appears to be sufficient to drive POA-AH^Gal^ pS6 expression nor is either behavior correlated with it, suggesting that POA-AH^Gal^ neurons are not driving egg care behaviors.

Although mating experience does alter parental care behavior and related medial POA-AH^Gal^ neuron activity in male mice (34), it is unlikely that the lack of POA-AH^Gal^ neuron activation in midshipman during egg care is explained by mating experience. In this experiment, we only used field-collected males from nests without eggs, suggesting those animals had not recently mated during the current reproductive season prior to their capture. However, during the experiment we included both males that had and males that had not mated in captivity in the egg care and control groups. All males show very low POA-AH^Gal^ activity, regardless of mating experience during the current season. Alternately, offspring developmental stage may play a factor. It is possible that POA-AH^Gal^ neurons are active during parental care directed towards larvae, which were not included in this study, rather than eggs. Midshipman males experience a shift in circulating androgen concentration when caring for larvae compared to caring for eggs (62). This shift could correspond with changes in POA-AH^Gal^ neuron activity during larval care, though further studies would be necessary to determine if this is the case. Nevertheless, the results of this experiment indicate that the large increase in POA-AH^Gal^ neuron activity seen during mating in courting males is not explained by the presence of eggs (i.e. early parental care).

An alternative explanation for our results is that POA-AH^Gal^ activity is related to sperm release. A population of neurons active during ejaculation in rats appears to receive galanin inputs (63), and electrical stimulation of the teleost POA-AH evokes sperm release (64). However, it is unlikely that sperm release alone explains the activity we find in POA-AH^Gal^ neurons of courting males for two reasons. First, mating satellite males of both morphs are also likely to be releasing sperm (4, 6) but do not show increased POA-AH^Gal^ activity. Second, in one mating trial no eggs were found in the nest when animals were collected, suggesting that although the courting male and female were in the nest together for the duration of the experiment, the female did not lay eggs and the male is unlikely to have released sperm, but still showed increased pS6 expression in its POA-AH^Gal^ neurons. Nevertheless, it is possible that some of the POA-AH^Gal^ activation we detect is related to sperm release.

Based on the results of the current study, we are unable to determine whether POA-AH^Gal^ activity seen during courting type I male mating results in galanin release, or if these neurons are regulating behavior through release of other peptides or transmitters. However, evidence from other studies suggests that galanin peptide is playing a role in male reproductive behavior. First, in midshipman, transcripts encoding galanin are upregulated in courting type I males during mating (43). Additionally, microinjection of galanin into the medial preoptic nucleus (a subdivision of the POA) of rats facilitates sexual behaviors (36, 37). Together, these studies show that galanin is involved in the regulation of reproductive behavior and suggest that the activity of POA-AH^Gal^ neurons in the present study is related to galanin release.

In summary, by co-labeling POA-AH neurons for galanin and pS6, we show that a molecularly-defined population of neurons increases activity during mating in courting type I male midshipman specifically, and that these neurons are not activated during other mating-related behaviors, uncovering POA-AH^Gal^ neurons as a potential neural substrate for the evolution of ARTs. More broadly, this work demonstrates that POA-AH^Gal^ neurons have a shared role across distant vertebrate lineages in social behaviors that directly contribute to individual fitness, including male mating in fish and mammals (38, 39) and mammalian parental care (39, 40), and provides functional evidence that allows for a better understanding of the evolutionary relationship between molecularly identified cell groups across both fish and mammals.

## Methods

### Animal subjects

Adult midshipman were collected from nests in California and Washington in May-August of 2016-2018 (65, 66). Each morph has distinguishing external characteristics including relative size and coloration that aid their recognition in the field; morph type was later confirmed on the basis of gonad type/size and swim bladder with attached sonic muscle size and morphology (6, 67). Fish were shipped overnight to Cornell University and housed in various sized aquaria (see below) with artificial seawater in environmental control rooms at 15-16°C with a 15:9 Light:Dark schedule. Vocal behavior was monitored using hydrophones (Aquarian Audio H1a, Anacordes, WA) as previously described (51). Behavior during mating, egg care, and nest defense experiments was recorded using video cameras (Canon Vixia HFR500) under red light. Analysis of videos was done using BORIS (68). Animal procedures were approved by the Institutional Animal Care and Use Committee of Cornell University.

### Mating behavior experiment

Males were held in large (100-200 gallon) aquaria divided into segments by plastic mesh (Fig. 2A). Each segment was approximately 1m × 1m and contained one large type I male (83.6-258.7 g, body weight; 17.8-26.0 cm, standard length), one small type I male (38.8-140.1 g, 14.2-21.4 cm), one type II male (6.5-18.2 g, 7.9-11.9 cm), and a single artificial nest made of a ceramic plate resting on a rim of bricks. Following the experimental design of prior studies, type I males were paired such that the larger male in the segment was expected to assume nest ownership and begin courting, while the smaller type I male would pursue the alternative tactic of cuckoldry (4, 5, 43).

Females (21.0-45.3 g, 11.8-14.8 cm) were added to a pair of neighboring nest segments 30 mins prior to the onset of dark (Fig. 2A), when courtship humming and mating occur (6, 51). Fish activity was observed by the experimenters under red light. Once a female entered the nest of either courting male, the nest in the neighboring segment was covered with a plastic mesh cage to prevent mating. Following an earlier study using the immediate early gene c-fos to identify neural activity in midshipman (69), fish were allowed to mate for two hours following female entry into a nest. The mating type I male and female along with neighboring non-mating control type I (i.e. male in nest covered by mesh cage) and non-mating control female were sacrificed two hours after the mating female entered the nest, and cuckolding type I and II males were sacrificed along with their neighboring control animals (i.e. type I and II males blocked from accessing a nest) two hours after they began cuckolding (Fig. 2A).

### Courtship humming behavior experiment

Type I males were housed in aquarium segments as described above (Fig. 3A). Each segment contained a single nest with one type I male. Vocal behavior was continuously monitored with hydrophones as previously described (51). Male pairs were monitored remotely using TeamViewer (v12.0.78517) and Audacity (v2.1.3) beginning at the start of the dark period. Males that hummed (48.87-210.53g, 15.5-25.0cm) at least 10 mins were collected two hours after the onset of humming (Fig 3A). Control males (36.18-207.36g, 15.9-24.4cm) were non-humming neighbors of humming males and were collected and sacrificed in parallel with humming males. Hydrophone recordings confirmed that control males did not vocalize during the two-hour experiment. Total humming duration was quantified using Raven Pro 1.5 (Cornell Lab of Ornithology) following previously described methods (51).

### Parental care behavior experiment

To generate stimulus nests with fertilized eggs, type I males were housed singly in aquaria (61cm × 56cm × 30.5cm) containing one artificial nest each. Gravid females were added to aquaria and allowed to mate. Parental males and females were removed after mating and replaced by experimental type I males. Males that mated were tested in unfamiliar nests, except in one case where the nest-holding male was tested in its own nest after being removed from the aquarium and then returned 24 hours later to determine whether POA-AH^Gal^ activity differed between males in nests with eggs they fertilized compared to males in nests with eggs fertilized by other males. Experimental males were added to aquaria containing nests that either had fertilized eggs (54.62-155.80g, 16.7-23.9cm) or no eggs (67.25-117.64g, 17.4-21.2cm), then were allowed to establish residency in nests (Fig. 4A). We expected that experimental type I males would take over nests and provide care given that nest takeover by type I males has been observed in captivity (6) and is common in nature (49, 70), and type I males will provide care for eggs that are not their own (61). Nesting males were sacrificed two hours after onset of the dark period, when midshipman are most active (51). Behaviors in the nest during the experiment were video recorded.

### Nest defense behavior experiment

To test whether pS6 expression in POA-AH^Gal^ neurons of courting type I males was dependent on nest defense against attempted cuckolders, we allowed courting type I males to mate either in the presence or absence of cuckolding males. Type I males were held singly in aquaria (see parental care behavioral experiment above) with one artificial nest. Males that mated without cuckolders (69.21-70.76g, each 17.5cm) were housed alone, while males that mated with cuckolders (43.06-108.65g, 15.1-20.1cm) were housed with one smaller type I male and one type II male. Once the courting male began humming, one or two females were added to the nest, and mating behavior was observed. Courting type I males were collected and sacrificed two hours after a female entered and remained in the nest (Fig. 5A).

### Immunohistochemistry

Directly following removal from tanks, animals were deeply anesthetized in 0.025% benzocaine (Sigma Aldrich, St. Louis, MO), then perfused with ice cold teleost ringer’s solution followed by ice cold 4% paraformaldehyde in 0.1M phosphate buffer (PB). Following perfusion, brains were removed and post-fixed in 4% paraformaldehyde for one hour at 4°C, then transferred to 0.1M PB for storage at 4°C. Brains were cryoprotected in 25% sucrose in 0.1M PB overnight at 4°C, then frozen in Tissue Plus O.C.T. compound (TissueTek, Torrance, CA) at −80°C. Brains were sectioned at 25μM in three series. Tissue sections were thaw mounted onto Superfrost Plus slides (ThermoFisher Scientific, Waltham, MA), allowed to dry overnight at room temperature, then were stored at −80°C.

IHC was performed on brain tissues collected from behaving (e.g. mating, humming) animals and controls in parallel. Prior to labelling, slides containing tissue sections were returned to room temperature and allowed to dry. Slides were washed three times for 10 min in phosphate buffered saline (PBS), followed by a 2 hour incubation in blocking solution of 0.2% bovine serum albumen (Sigma Aldrich), 0.3% Triton-X 100 (Sigma Aldrich), and 10% normal goat serum (NGS, ThermoFisher) in PBS, then 18 hours in guinea pig anti-galanin (1:250, custom raised against midshipman galanin peptide; Pocono Rabbit Farm and Laboratory, Canadensis, PA) and rabbit anti-pS6 (1:250, Cell Signaling Technology, Danvers, MA, Cat# 4858, RRID:AB_916156) primary antibodies in blocking solution at room temperature. Following primary antibody incubation, slides were washed three times in PBS, then incubated 2 hours in goat anti-guinea pig secondary antibody conjugated to Alexa Fluor 488 (1:500, Life Technologies, Carlsbad, CA, Cat# A-11073, RRID:AB_142018) and donkey anti-rabbit secondary antibody conjugated to Alexa Fluor 568 (1:200, Life Technologies, Cat# A10042, RRID:AB_2534017) in PBS+10% NGS. After secondary antibody incubation, slides were washed in PBS three times for 10 min, followed by one 10 min wash in double distilled water, and then coverslipped with ProLong Gold with DAPI (ThermoFisher). After coverslipping, slides were allowed to dry at room temperature overnight, then edges were sealed with nail polish. Slides were stored at 4°C. Specificity of anti-galanin antibody was confirmed by performing IHC following preadsorption of antibody with 50μM galanin peptide (Pocono) or omission of primary antibody. A prior study has demonstrated specificity of anti-pS6 (71) primary antibody in teleost fish. Western blot confirmed anti-pS6 specificity for midshipman.

### Image acquisition and processing

All images through the POA-AH were acquired on a Zeiss LSM 880 confocal microscope (Cornell University Biotechnology Resource Center Imaging Facility, NIH S10OD018516). The POA-AH of each fish was imaged bilaterally at 20X with a 10-level z-stack (5μM optical section, 2.5μM step). Tile scanning with 20% overlap and medium (0.70) stitching threshold was used when necessary to capture the entire POA-AH in an image. Quantification of pS6-expressing galanin neurons was done using FIJI (72). Green and red channels of each image were merged, then z-stacks were projected using the maximum intensity function. Cell bodies of POA-AH^Gal^ neurons were first identified using the following criteria: 1) a clearly defined perimeter and 2) an identifiable nucleus and/or neurite; observers then determined whether the POA-AH^Gal^ neuron expressed pS6. Criteria for pS6 expression in POA-AH^Gal^ neurons were either visually identifiable overlap of green and red signal (i.e. yellow signal) or red signal visible within the perimeter of the green POA-AH^Gal^ cell body. Counting of POA-AH^Gal^ neurons expressing pS6 was performed by an observer blind to the condition of each animal. Images were cropped and re-sized for figures using Adobe Photoshop CS6 (Adobe Systems Incorporated, San Jose, CA).

### Statistics

Statistical tests were conducted using R (3.3.3). Comparisons of number of pS6-expressing POA-AH^Gal^ neurons in mating experiment were made by one-way ANOVA and post hoc comparisons made with Tukey’s HSD test. Comparison of number of pS6-expressing POA-AH^Gal^ neurons in courtship humming experiment was made using a Welch two sample t-test to correct for unequal variance between groups. Comparison of number pS6-expressing POA-AH^Gal^ neurons during egg care was made using an two sample t-test. Correlations of humming and parenting behaviors with pS6 expression in POA-AH^Gal^ neurons were tested using Pearson’s correlation coefficient. Due to limited sample size, only descriptive statistics were used to describe data from nest defense experiment.

## ACKNOWLEDGEMENTS

We thank Clara Liao, Margaret Marchaterre, Rich Moore, Kevin Rohmann, David Rose, Elisabeth Rosner, and Rose Tatarsky for technical assistance; Joe Sisneros for assistance obtaining collection permits; Karen Maruska for providing a protocol for pS6 immunohistochemistry; Irene Ballagh, Eric Schuppe, and Rose Tatarsky for constructive feedback on drafts of the manuscript; and the Cornell University Statistical Consulting Unit for advice on analyses. This work was generously supported by NSF IOS 1656664, Cornell University, and the Department of Neurobiology and Behavior.

## AUTHOR CONTRIBUTIONS

J.A.T. and A.H.B. conceived and designed the study and wrote the manuscript. J.A.T., I.S., A.M., and A.H.B. conducted the experiments and analyzed the data. All authors approved the final manuscript.

## DECLARATION OF INTERESTS

The authors declare no competing interests

## References

1. Godwin J (2010) Neuroendocrinology of sexual plasticity in teleost fishes. Front Neuroendocrinol 31(2):203–16.

2. Feng NY, Bass AH (2017) Neural, hormonal, and genetic mechanisms of alternative reproductive tactics: Vocal fish as model systems. Hormones, Brain, and Behavior, eds Pfaff DW, Joels M, pp 47–68. 3rd Ed.

3. Mank JE, Avise JC (2006) Comparative phylogenetic analysis of male alternative reproductive tactics in ray-finned fishes. Evolution 60(6):1311–1316.

4. Lee JSF, Bass AH (2004) Does exaggerated morphology preclude plasticity to cuckoldry in the midshipman fish (*Porichthys notatus*)? Naturwissenschaften 91(7):338–41.

5. Lee JSF, Bass AH (2006) Dimorphic male midshipman fish: Reduced sexual selection or sexual selection for reduced characters? Behav Ecol 17(4):670–675.

6. Brantley RK, Bass AH (1994) Alternative male spawning tactics and acoustic signals in the plainfin midshipman fish *Porichthys notatus* Girard (Teleostei, Batrachoididae). Ethology 96:213–232.

7. Arora HL (1948) Observations on the habits and early life history of the Batrachoid fish, *Porichthys notatus* Girard. Copeia 1948(2):89–93.

8. McKibben JR, Bass AH (1998) Behavioral assessment of acoustic parameters relevant to signal recognition and preference in a vocal fish. J Acoust Soc Am 104(6):3520–33.

9. Brantley RK, Wingfield JC, Bass AH (1993) Sex steroid levels in *Porichthys notatus*, a fish with alternative reproductive tactics, and a review of the hormonal bases for male dimorphism among teleost fishes. Horm Behav 27(3):332–47.

10. Arterbery AS, Deitcher DL, Bass AH (2010) Divergent expression of 11beta-hydroxysteroid dehydrogenase and 11beta-hydroxylase genes between male morphs in the central nervous system, sonic muscle and testis of a vocal fish. Gen Comp Endocrinol 167(1):44–50.

11. Bass AH (1992) Dimorphic male brains and alternative reproductive tactics in a vocalizing fish. Trends Neurosci 15(1987):139–145.

12. Grober MS, Fox SH, Laughlin C, Bass AH (1994) GnRH cell size and number in a teleost fish with two male reproductive morphs: Sexual maturation, final sexual statues and body size allometry. Brain Behav Evol 43:61–78.

13. Bass AH, Horvath BJ, Brothers EB (1996) Nonsequential developmental trajectories lead to dimorphic vocal circuitry for males with alternative reproductive tactics. J Neurobiol 30(4):493–504.

14. Bass AH, Baker R (1990) Sexual dimorphisms in the vocal control system of a teleost fish: Morphology of physiologically identified neurons. J Neurobiol 21(8):1155–1168.

15. Goodson JL, Bass AH (2000) Forebrain peptides modulate sexually polymorphic vocal circuitry. Nature 403:769–772.

16. Remage-Healey L, Bass AH (2004) Rapid, hierarchical modulation of vocal patterning by steroid hormones. J Neurosci 24(26):5892–900.

17. Remage-Healey L, Bass AH (2007) Plasticity in brain sexuality is revealed by the rapid actions of steroid hormones. J Neurosci 27(5):1114–22.

18. Küpper C, et al. (2015) A supergene determines highly divergent male reproductive morphs in the ruff. Nat Genet 48(1):79–83.

19. Lamichhaney S, et al. (2015) Structural genomic changes underlie alternative reproductive strategies in the ruff (*Philomachus pugnax*). Nat Genet 48(1):84–88.

20. Zamudio KR, Sinervo B (2000) Polygyny, mate-guarding, and posthumous fertilization as alternative male mating strategies. Proc Natl Acad Sci 97(26):14427–14432.

21. Maggioncalda AN, Sapolsky RM (2002) Disturbing behaviors of the orangutan. Sci Am 60–65. Available at: http://www.nature.com/scientificamericanmind/journal/v20/n3/full/scientificamericanmind0509-14.html [Accessed November 13, 2014].

22. Goodson JL, Bass AH (2001) Social behavior functions and related anatomical characteristics of vasotocin/vasopressin systems in vertebrates. Brain Res Rev 35(3):246–65s.

23. Godwin J, Thompson R (2012) Nonapeptides and social behavior in fishes. Horm Behav 61(3):230–8.

24. Phelps SM, Okhovat M, Berrio A (2017) Individual differences in social behavior and cortical vasopressin receptor: Genetics, epigenetics, and evolution. Front Neurosci 11(OCT):1–12.

25. Bass AH, Grober MS (2001) Social and neural modulation of sexual plasticity in teleost fish. Brain, Behav Evol 57:293–300.

26. Goodson JL, Bass AH (2000) Vasotocin innervation and modulation of vocal-acoustic circuitry in the teleost *Porichthys notatus*. J Comp Neurol 422(3):363–79.

27. Donaldson ZR, Young LJ (2008) Oxytocin, vasopressin, and the neurogenetics of behavior. Science 322. doi:10.1126/science.1158668.

28. Kelly AM, Goodson JL (2014) Social functions of individual vasopressin-oxytocin cell groups in vertebrates: What do we really know? Front Neuroendocrinol 35(4):512–529.

29. Koshiyama H, et al. (1987) Central galanin stimulates pituitary prolactin secretion in rats: Possible involvement of hypothalamic vasoactive intestinal polypeptide. Neurosci Lett 75(1):49–54.

30. Murakami Y, et al. (1987) Galanin stimulates growth hormone (GH) secretion via GH-releasing factor (GRF) in conscious rats. Eur J Pharmacol 136(3):415–418.

31. Sahu A, Crowley WR, Tatemoto K, Balasubramaniam A, Kalra SP (1987) Effects of neuropeptide Y, NPY analog (norleucine4-NPY), galanin and neuropeptide K on LH release in ovariectomized (ovx) and ovx estrogen, progesterone-treated rats. Peptides 8(5):921–926.

32. Kyrkouli S, Stanley BG, Leibowitz SF (1986) Galanin: Stimulation of feeding induced by medial hypothalamic injection of this novel peptide. Eur J Pharmacol 122:159–160.

33. Qualls-Creekmore E, et al. (2017) Galanin-expressing GABA neurons in the lateral hypothalamus modulate food reward and non-compulsive locomotion. J Neurosci 37(25):0155–17.

34. Dulac CG, O’Connell LA, Wu Z (2014) Neural control of maternal and paternal behaviors. Science 345(6198):765–770.

35. Fischer EK, O’Connell LA (2017) Modification of feeding circuits in the evolution of social behavior. J Exp Biol 220(1):92–102.

36. Bloch GJ, Butler PC, Kohlert JG, Bloch DA (1993) Microinjection of galanin into the medial preoptic nucleus facilitates copulatory behavior in the male rat. Physiol Behav 54(4):615–624.

37. Bloch GJ, Butler PC, Kohlert JG (1996) Galanin microinjected into the medial preoptic nucleus facilitates female- and male-typical sexual behaviors in the female rat. Physiol Behav 59(6):1147–1154.

38. Bakker J, Woodley SK, Kelliher KR, Baum MJ (2002) Sexually dimorphic activation of galanin neurones in the ferret’s dorsomedial preoptic area/anterior hypothalamus after mating. J Neuroendocrinol 14(2):116–125.

39. Wu Z, Autry AE, Bergan JF, Watabe-Uchida M, Dulac CG (2014) Galanin neurons in the medial preoptic area govern parental behaviour. Nature 509(7500):325–30.

40. Kohl J, et al. (2018) Functional circuit architecture underlying parental behaviour. Nature 556(7701):326–331.

41. Moffitt JR, et al. (2018) Molecular, spatial, and functional single-cell profiling of the hypothalamic preoptic region. Science 5324(November):1–22.

42. Partridge CG, MacManes MD, Knapp R, Neff BD (2016) Brain transcriptional profiles of male alternative reproductive tactics and females in bluegill sunfish. PLoS One 11(12):1–21.

43. Tripp JA, Feng NY, Bass AH (2018) Behavioural tactic predicts preoptic-hypothalamic gene expression more strongly than developmental morph in fish with alternative reproductive tactics. Proc R Soc B 285(1871).

44. Bengston SE, et al. (2018) Genomic tools for behavioural ecologists to understand repeatable individual differences in behaviour. Nat Ecol Evol 2(6):944–955.

45. Knight ZA, et al. (2012) Molecular profiling of activated neurons by phosphorylated ribosome capture. Cell 151(5):1126–1137.

46. Foran CM, Myers DA, Bass AH (1997) Modification of gonadotropin releasing hormone (GnRH) mRNA expression in the retinal-recipient thalamus. Gen Comp Endocrinol 106(2):251–64.

47. Foran CM, Bass AH (1998) Preoptic AVT immunoreactive neurons of a teleost fish with alternative reproductive tactics. Gen Comp Endocrinol 111(3):271–82.

48. Zeddies DG, Fay RR, Alderks PW, Shaub KS, Sisneros JA (2010) Sound source localization by the plainfin midshipman fish, *Porichthys notatus*. J Acoust Soc Am 127(5):3104–13.

49. Bose APH, Mcclelland GB, Balshine S (2016) Cannibalism, competition, and costly care in the plainfin midshipman fish, *Porichthys notatus*. Behav Ecol 27(2):628–636.

50. Sisneros JA, Alderks PW, Leon K, Sniffen B (2009) Morphometric changes associated with the reproductive cycle and behaviour of the intertidal-nesting, male plainfin midshipman *Porichthys notatus*. J Fish Biol 74(1):18–36.

51. Feng NY, Bass AH (2016) “Singing” fish rely on circadian rhythm and melatonin for the timing of nocturnal courtship vocalization. Curr Biol 26(19):2681–2689.

52. Sisneros JA, Forlano PM, Knapp R, Bass AH (2004) Seasonal variation of steroid hormone levels in an intertidal-nesting fish, the vocal plainfin midshipman. Gen Comp Endocrinol 136(1):101–16.

53. Renn SCP, Aubin-Horth N, Hofmann HA (2008) Fish and chips: Functional genomics of social plasticity in an African cichlid fish. J Exp Biol 211(Pt 18):3041–56.

54. Hokfelt T ed. (2010) Galanin (Springer, New York).

55. Puelles L, Harrison M, Paxinos G, Watson C (2013) A developmental ontology for the mammalian brain based on the prosomeric model. Trends Neurosci 36(10):570–578.

56. Domínguez L, Morona R, González A, Moreno N (2013) Characterization of the hypothalamus of *Xenopus laevis* during development. I. The alar regions. J Comp Neurol 521(4):725–759.

57. Affaticati P, et al. (2015) Identification of the optic recess region as a morphogenetic entity in the zebrafish forebrain. Sci Rep 5:8738.

58. Bass AH, Forlano PM (2008) Neuroendocrine mechanisms of alternative reproductive tactics: The chemical language of reproductive and social plasticity. Alternative Reproductive Tactics: An Integrative Approach, eds Oliveira RF, Taborsky M, Brockman HJ (Cambridge University Press), pp 109–131.

59. Herget U, Wolf A, Wullimann MF, Ryu S (2014) Molecular neuroanatomy and chemoarchitecture of the neurosecretory preoptic-hypothalamic area in zebrafish larvae. J Comp Neurol 522(7):1542–1564.

60. Forlano PM, Cone RD (2007) Conserved neurochemical pathways involved in hypothalamic control of energy homeostasis. J Comp Neurol 505:235–248.

61. Bose APH, Kou HH, Balshine S (2016) Impacts of direct and indirect paternity cues on paternal care in a singing toadfish. Behav Ecol 27(5):1507–1514.

62. Knapp R, Wingfield JC, Bass AH (1999) Steroid hormones and paternal care in the plainfin midshipman fish (*Porichthys notatus*). Horm Behav 35(1):81–9.

63. Veening JG, Coolen LM (1998) Neural activation following sexual behavior in the male and female rat brain. Behav Brain Res 92(2):181–93.

64. Demski LS, Bauer DH, Gerald JW (1975) Sperm release evoked by electrical stimulation of the fish brain: A functional-anatomical study. J Exp Zool 191(2):215–232.

65. Bass AH (1996) Shaping brain sexuality. Am Sci 84(4):352–363.

66. McIver EL, Marchaterre MA, Rice AN, Bass AH (2014) Novel underwater soundscape: Acoustic repertoire of plainfin midshipman fish. J Exp Biol 217:2377–2389.

67. Bass AH, Marchaterre MA (1989) Sound-generating (sonic) motor system in a teleost fish (*Porichthys notatus*): Sexual polymorphisms and general synaptology of sonic motor nucleus. J Comp Neurol 286:141–153.

68. Friard O, Gamba M (2015) Behavioral Observation Research Interactive Software (BORIS). Available at: http://penelope.unito.it/boris?page=home%5Cnhttp://www.boris.unito.it/?page=home.

69. Petersen CL, et al. (2013) Exposure to advertisement calls of reproductive competitors activates vocal-acoustic and catecholaminergic neurons in the plainfin midshipman fish, *Porichthys notatus*. PLoS One 8(8):e70474.

70. Cogliati KM, Neff BD, Balshine S (2013) High degree of paternity loss in a species with alternative reproductive tactics. Behav Ecol Sociobiol 67(3):399–408.

71. Butler JM, Whitlow SM, Roberts DA, Maruska KP (2018) Neural and behavioural correlates of repeated social defeat. Sci Rep 8(1):1–13.

72. Schindelin J, et al. (2012) Fiji: An open source platform for biological image analysis. Nat Methods 9(7):676–682.

