## Supporting Information for "Preoptic galanin neuron activation is specific to courtship reproductive tactic in fish with two male morphs"

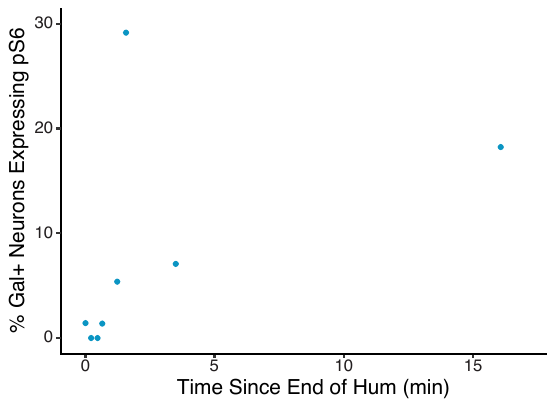

**Figure S1.** Relates to Figure 2. Proportion of POA-AH^Gal^ neurons expressing pS6 is not correlated with length of time between end of humming and removal from nest for humming type I males. Pearson’s correlation p=0.2451, r=0.465508.

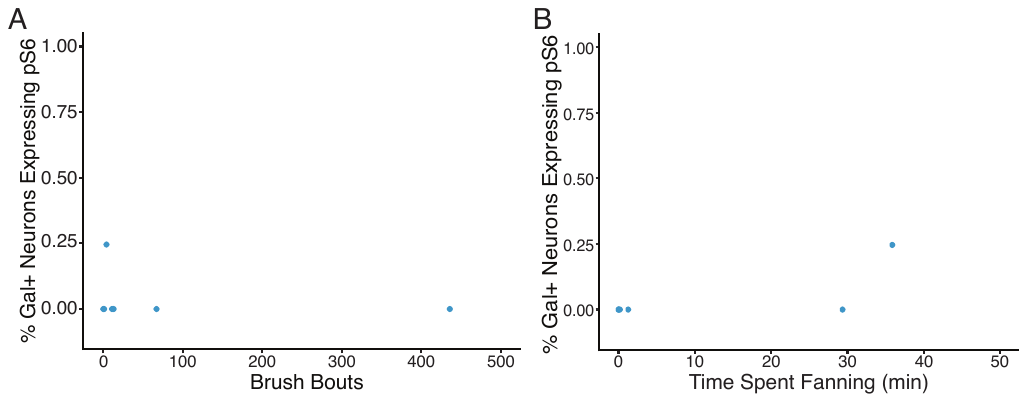

**Figure S2.** Relates to Figure 3. Proportion of POA-AH^Gal^ neurons expressing pS6 is not correlated with A) the number of bouts of brushing eggs with mouth, Pearson’s correlation p=0.6706, r=-0.1978782 or B) the amount of time spent fanning fins in the nest, Pearson’s correlation p=0.06185, r=0.7312215

**Table S1.** Midshipman morph differences

| **Trait** | **Type I male** | **Type II male** | **Female** |
| --- | --- | --- | --- |
| Behavioral |  |  |  |
| Nest building | Yes | No | No |
| Egg guarding | Yes | No | No |
| Vocal repertoire | Hums, grunts, grunt trains, growls | Grunts | Grunts |
| Call duration | Long | Short | Short |
| Call fundamental frequency | High | Low | Low |
| Somatic |  |  |  |
| Body size | Large | Small | Intermediate |
| Gonad/body size ratio | Small | Large | Large (gravid); small (spent) |
| Ventral coloration | Olive-gray | Mottled yellow | Bronze/golden (gravid); mottled (spent) |
| Vocal motor |  |  |  |
| Vocal muscle mass, fiber number, and diameter | Large | Small | Small |
| Vocal motor neuron size | Large | Small | Small |
| Endocrine |  |  |  |
| Predominant circulating steroids | 11-ketotestosterone | Testosterone | Testosterone, estradiol |
| Neuroendocrine |  |  |  |
| Aromatase activity, mRNA, protein | Low | High | High |
| GnRH-POA neuron size | Large | Small | Small |
| GnRH-POA neuron number/body size ratio | Low | High | Low |
| AVT-POA neuron size | Large | Small | Large |
| AVT-POA neuron number/body size ratio | Low | High | Low |
| vocal-motor circuit response to exogenous hormones |  |  |  |
| 11-ketotestosterone | Facilitation | No effect | No effect |
| Testosterone | No effect | AR-dependent facilitation | ER-dependent facilitation |
| Estradiol | Facilitation | Facilitation | Facilitation |
| Cortisol | Facilitation | Suppression | Suppression |
| Arginine vasotocin | Suppression | No effect | No effect |
| Isotocin | No effect | Suppression | Suppression |

Table adapted from Feng & Bass, 2017 & Tripp, Feng, & Bass 2018.

**Movie 1.** Type I male courtship tactic. Courting type I male excavates gravel from interior of artificial nest in laboratory aquarium, hums, and grabs approaching female. Recorded under red light during dark period using a video camera (Canon) with attached hydrophone (Aquarian Audio).

**Movie 2.** Satellite cuckolding tactic. Overhead view of cuckolding type I male satellite mating at a nest containing courting type I male and female. Satellite male with its tail inserted into the nest performs a spawning reflex. Recorded under red light during dark period using an iPhone 6.
